# *Aedes vittatus* in Haiti: continued expansion across the Caribbean and implications for the Americas

**DOI:** 10.64898/2026.06.15.732314

**Authors:** Daphenide St-Louis, Philippe Boussès, Loîc Talignani, Chrystelle Lasica, Evens Emmanuel, Fréderic Simard, David Roiz

## Abstract

**Background:** *Aedes vittatus*, an emerging invasive species and arboviral vector with high ecological plasticity and dispersal capacity, is expanding in the Caribbean, being detected in the Dominican Republic, Cuba, and Jamaica and recently in Yucatán, Mexico.

**Objectives:** This study aimed to provide evidence of the establishment of *Ae. vittatus* in Haiti.

**Methods:** Larval surveys were done across 12 localities around Cap-Haïtien, in the North-East department of Haiti, the specimens were identified morphologically and confirmed by COI DNA barcoding.

**Findings:** We report, for the first time, the detection of the invasive vector *Aedes (Fredwardsius) vittatus* in Haiti, confirmed morphologically and molecularly, and being established in half of the municipalities sampled. Phylogenetic analyses group Haitian specimens with Caribbean and American populations, closely related to Mexico, indicating regional expansion.

**Main conclusion:** Given its role as a vector of chikungunya, dengue, Zika and yellow fever, strengthened entomological surveillance and early detection in North America and the Caribbean are urgently needed.

## Introduction

Invasive vector mosquito species, such as *Aedes aegypti* and *Aedes albopictus*, have spread globally through human activities increasing the public health and economic burden of arboviral diseases such as dengue, chikungunya and Zika^(1,2)^. *Aedes vittatus*, an emerging invasive species and arboviral vector with high ecological plasticity and dispersal capacity, was first detected in the Caribbean in 2019 in the Dominican Republic and Cuba, followed by Jamaica in 2023 and Mexico in 2024–2025^(3,4,5)^. These records indicate rapid expansion across the Caribbean basin and into the American continent. Despite demonstrated vector competence, the species remains poorly studied^(6)^. Haiti, characterized by high population density, environmental change, and intense human mobility, provides favorable conditions for mosquito invasions; however, *Ae. vittatus* had not previously been reported there. We report the first detection and evidence of establishment of *Ae. vittatus* in Haiti.

## Materials and methods

### Study Area

The survey was conducted in the North and North-East departments of Haiti, restricted to the extensive low-elevation coastal zones of both regions. Following the 2010 earthquake that struck the Port-au-Prince area, the North department experienced the arrival of more than 2.3 million people, resulting in rapid and largely unplanned urban expansion^6^. This department is therefore highly populated (500 inhabitants/km^2^), in contrast to the North-East, where population density ranges around 240 inhabitants/km^2^.

The climate varies from tropical monsoon (North Department) to tropical savanna (North East Department), with mean annual temperatures of 24 to 28 °C and seasonal rainfall (April to November), averaging between 1,000 and 1,500 mm per year, with a decreasing gradient from the coast toward the interior.

The study area, located mainly within the coastal plains surrounding the departmental capital, Cap-Haïtien, extended from Limbé (19°42’N, 72°24’W) in the west to Ouanaminthe (19°29’N, 71°46’W), located on the Dominican border 75 km to the east, following the coastal fringe and reaching up to 20 km inland. It is characterized by moderately low relief, most of the coastline and its surroundings lie at elevations generally below 100 m in urban and peri-urban zones.

### Mosquito sampling

Between October 7 and October 31, 2025, during the latter part of the rainy season, surveys were conducted in urban, peri-urban and rural areas across both departments. A total of 23 larval habitats were collected in 18 localities within 12 communes (7 in the North and 5 in the North East) (Figure 1). Sampling time did not exceed 3 hours per commune. Mosquito larvae were collected from peridomestic and public spaces. For each sample site, some collected larvae were mounted on slide and a part was reared to the adult stage, then both were deposited in the ARIM-IRD collection^(7)^. Geographical coordinates and data were uploaded to the GBIF platform. This study was conducted with authorization from the Haitian Ministry of Environment, and mosquito importation from Haiti to France was approved by the Departmental Directorate for Population Protection (DDPP).

**Figure 1.**
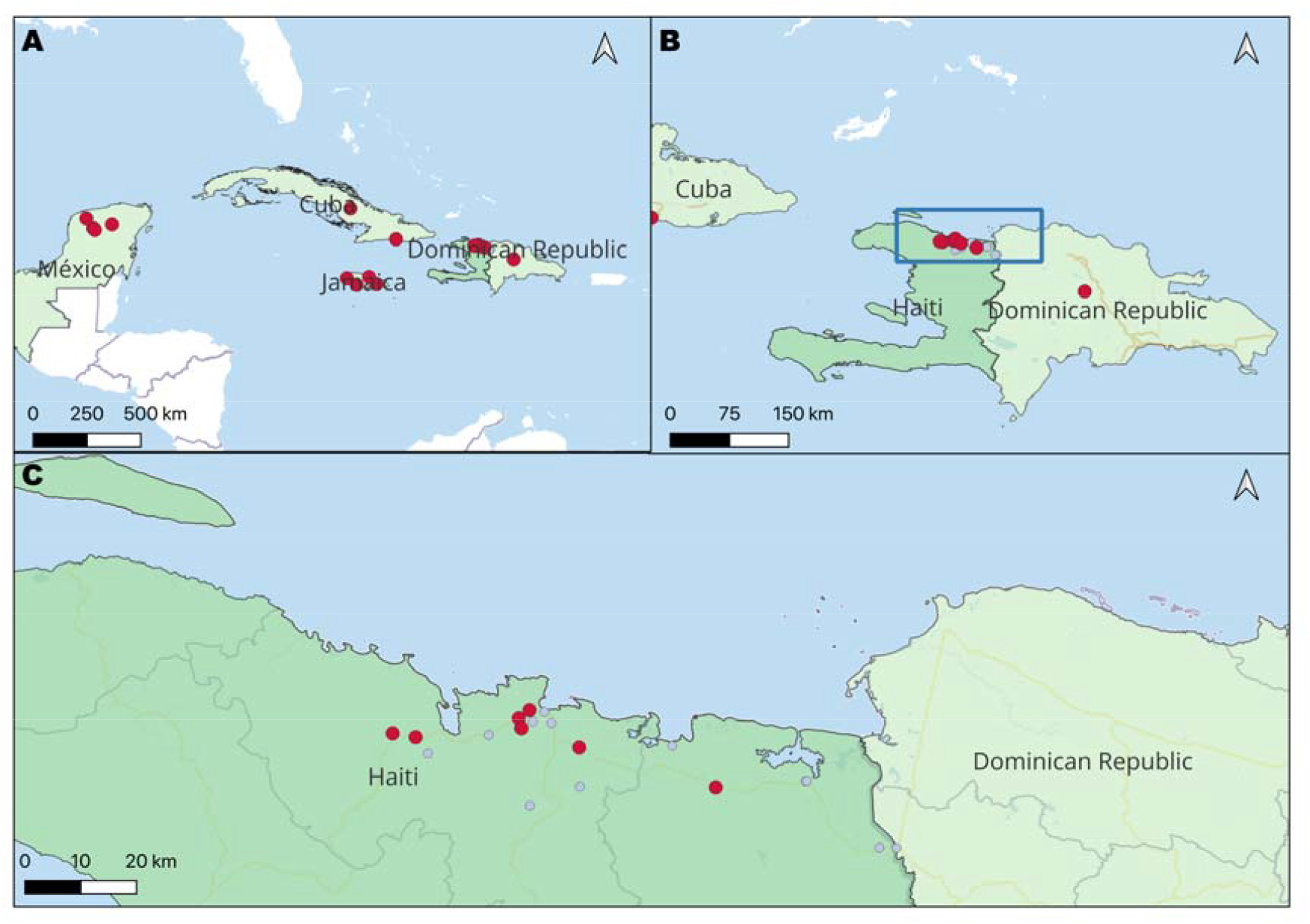
Location of the sampling sites. A. Situation map of Haiti within Caribbean countries with presence (green area) and location of capture points (red dots) of *Aedes vittatus* in neighboring countries; B. Delimitation of study area in Haiti (blue rectangle) and location capture (red dots) of *Ae. vittatus* in Haiti and the Dominican Republic; C. Location of the breeding site of *Ae. vittatus* in the study area (red dots) and other collection sites negative to *Ae. vittatus* (grey dots) (Table I).

### Morphological and molecular identification with COI-based DNA barcoding

The invasive species *Ae. vittatus* was identified using European^(8)^, African^(9)^ and Sahelian^(10)^ taxonomic keys, and confirmed with the species redescription of Huang^(11)^. Five samples from the localities Quartier Morin (2 adults: QUA003-Haiti06 and 07) and Lory, Milot (2 adults: MIL001-Haiti08 and 09 and 1 larvae: MIL004LAV-10) were used for molecular identification. Mosquito DNA was extracted with the Qiagen Blood&tissue kit (Qiagen, Valencia, USA). Mosquitoes were individually crushed with 3 metal beads in a Qiagen TissueLyserII at 30Hz for 3 minutes. The rest of the protocol followed the manufacturer’s instructions. PCR was performed in duplicate of these samples to validate the species identification through a universal invertebrate-specific primer pair LCO1490 (5′-GGT CAA CAA ATC ATA AAG ATA TTG G-3′) and HCO2198 (5′-TAA ACT TCA GGG TGA CCA AAA AAT CA-3′) to amplify a 658-bp fragment of the mitochondrial gene cytochrome c oxidase subunit I barcoding region (CO1) ^(12)^. One PCR reaction was made in a total volume of 35µL containing 17.5µL Invitrogen TM PlatinumTM Hot Start PCR 2X Master Mix, 0.35µL of each primer, 9.8µL of biology grade water, and 7µL of DNA. PCR amplification products were sent to Macrogen (Macrogen Europe, Amsterdam, the Netherlands) for sanger sequencing. A Basic Local Alignment Search Tool (BLAST) analysis was conducted in the National Center for Biotechnology Information (NCBI) GenBank database to explore species identification based on the amplified and sequenced molecular product^(13)^.

### Sequence assembly, alignment and Phylogenetic analyses

Forward and reverse reads of four samples were assembled and edited to generate consensus sequences using CodonCode Aligner v12.0.4 (https://www.codoncode.com/aligner). Sequence alignments were performed with MAFFT v7 via the online server (https://mafft.cbrc.jp/alignment/server/) using default parameters^(14)^. The resulting alignments were visually inspected and manually edited in AliView v1.28^(15)^. The best-fit nucleotide substitution model was determined using jModelTest2 through the CIPRES Science Gateway (https://www.phylo.org/) under default settings^(16)^. Mitochondrial sequences, including COX1 marker sequences of *Aedes spp*., were retrieved from GenBank (accession numbers provided in the phylogenetic tree). The dataset comprised sequences from Africa, Asia, Europe, the Caribbean, as well as newly generated sequence from Mexico.

Phylogenetic relationships among *Aedes vittatus* sequences were inferred using BEAST v1.10.5, applying a General Time Reversible substitution model+⍰returned by jModelTest2, a strict molecular clock, and a coalescent constant size tree prior. *Psorophora ferox* was designated as the outgroup^(17)^. Markov chain Monte Carlo (MCMC) analyses were run for 10 million generations, with trees sampled every 100 generations. Effective sample sizes (ESS) were assessed using Tracer v 1.7.2^(18)^, with values >200 considered adequate. Convergence and stationarity were assessed by examining parameter traces, and the first 25% of samples were discarded as burn-in prior to summarizing posterior probabilities. The final phylogenetic tree was visualized and edited using FigTree v1.4.4^(19)^.

### Network analysis

Analyses at the intraspecific level were performed on datasets that encompassed only *Ae. vittatus* sequences. The genetic differentiation between specimens belonging to the major phylogroups inferred by the phylogenetic analyses of the combined dataset was assessed for each COX1 gene using DnaSP v.6.12.03^(20)^. Haplotype networks were constructed using the median-joining method^(21)^, which has the ability to deal with missing data as well as to infer ancestral haplotypes. This method also performs well against or outperforms other network approaches^(22)^. It was implemented using the software PopART v1.7^(23)^, with epsilon value set to 0 in order to minimize alternative median networks. The resulting networks were visualized and edited in Inkscape v1.2.2 (https://inkscape.org/).

## Results

Larval surveys conducted during October 7–24, 2025 identified *Ae. vittatus* in 8 of 23 sites (34.7%) across six municipalities in northern Haiti (Cap-Haïtien, Quartier Morin, Milot, Limbé, Trou du Nord, and Terrier Rouge) (Table I). The species was found alone in three of these sites and co-occurred with *Ae. albopictus* (3 sites) and *Ae. aegypti* (2 sites). Breeding habitats included diverse peridomestic and public environments across urban, peri-urban, and rural settings, such as water storage containers, groundwater accumulations, poultry waterers, wastewater channels, puddles, and tree root cavities. These habitats were generally devoid of vegetation and varied in water volume and organic content. Eight additional mosquito species were recorded, including *Ae. aegypti, Ae. albopictus, Culex quinquefasciatus, Cx. garciai, Cx. interrogator, Cx. nigripalpus, Psorophora confinnis* and *Ps. pygmaea*. Note that *Cx. garciai* and *Cx. interrogator* constitute new country records for Haiti.

**Table 1.**
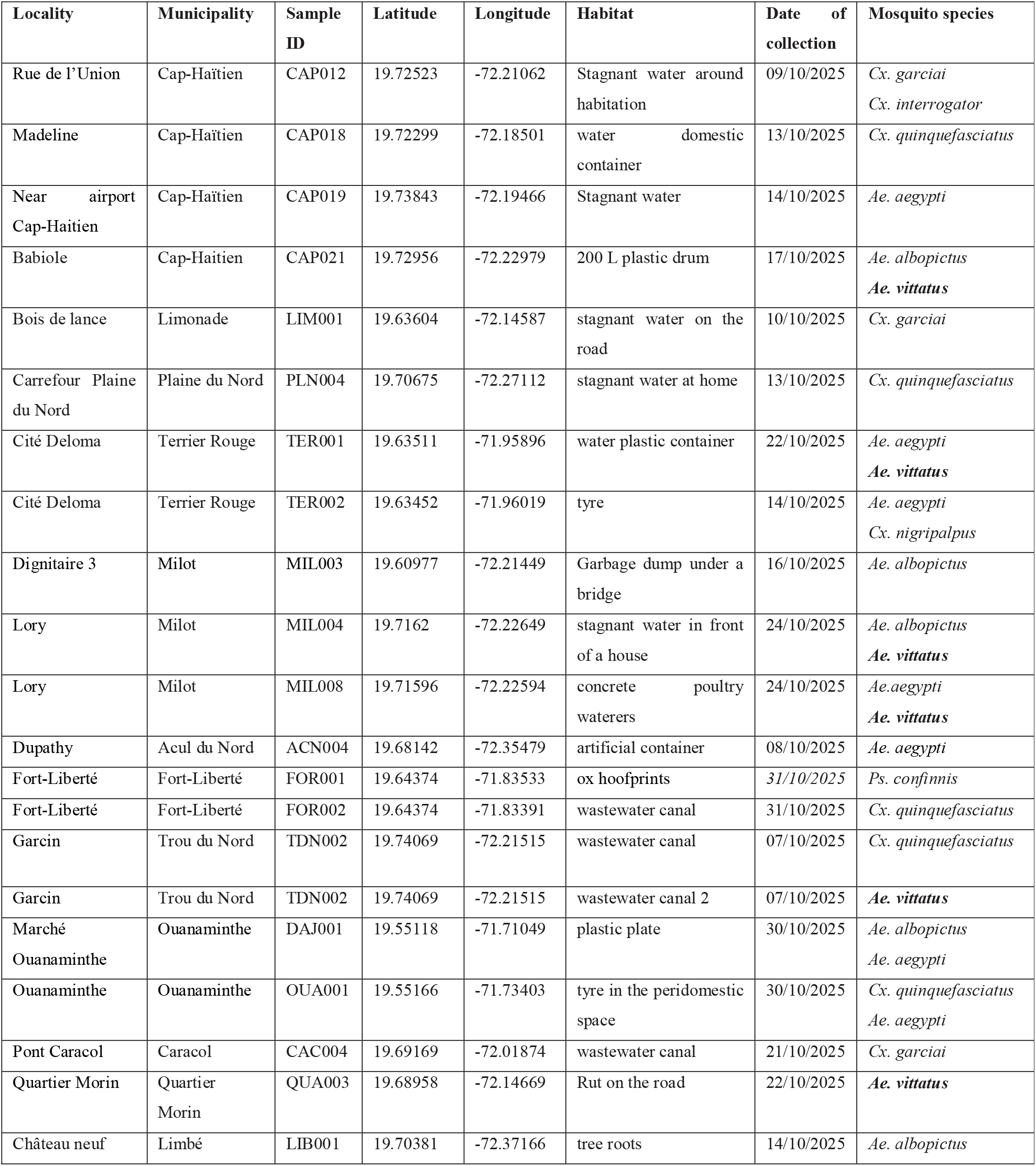
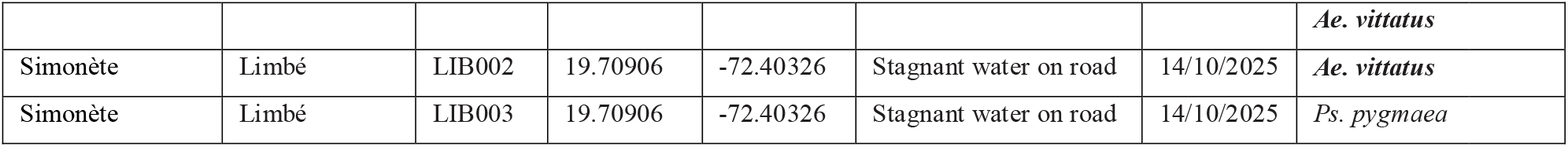
Samples and mosquito species collected in North and Northeast department of Haiti.

### Morphological identification

Both larval stages and adults obtained from breeding were examined, and diagnostic characters for the principal life stages for *Ae. vittatus* are summarized as follows (Figure 2). **Female**. Head with a proboscis dark with a median spattering of pale yellowish scales. <u>Thorax</u>. Scutum with acrostichal setae; 3 pairs of distinct, small, white spots of narrow scales on the anterior two-thirds of the scutum; scutellum with broad white scales on all three lobes; lower mesepimeral setae present; tibia with a median white ring, tarsomeres III-1-4 with large basal white bands, tarsomere III-5 all white. <u>Abdomen</u>. Tergite I with a large median white spot. **Male Genitalia**. Gonocoxite elongate, without basal and apical lobes. The gonostyle has a very characteristic shape, strongly enlarged and globose at its tip, with a long, strongly curved apical spine located at the base of the globose part. The claspette is large, with a narrow base, and distinctly swollen from its middle. **Larva** (Figure 2). <u>Head</u>. Setae 5-C single or sometimes double, 6-C single. <u>Abdomen</u>. Seta 6-I-IV usually with 2 long branches; pecten with 19-25 very closely spaced spines, with 3-4 basal denticles; only the most (rarely two) distal spine more widely spaced and implanted beyond seta 1-S; comb of 8 (6-10) scales in a single irregular row; saddle incomplete without marginal spicules; 1-X single, 2-X with 5 branches; 4 precratal setae; anal papillae long, about 4 times as long as saddle, slender and tapered.

**Figure 2.**
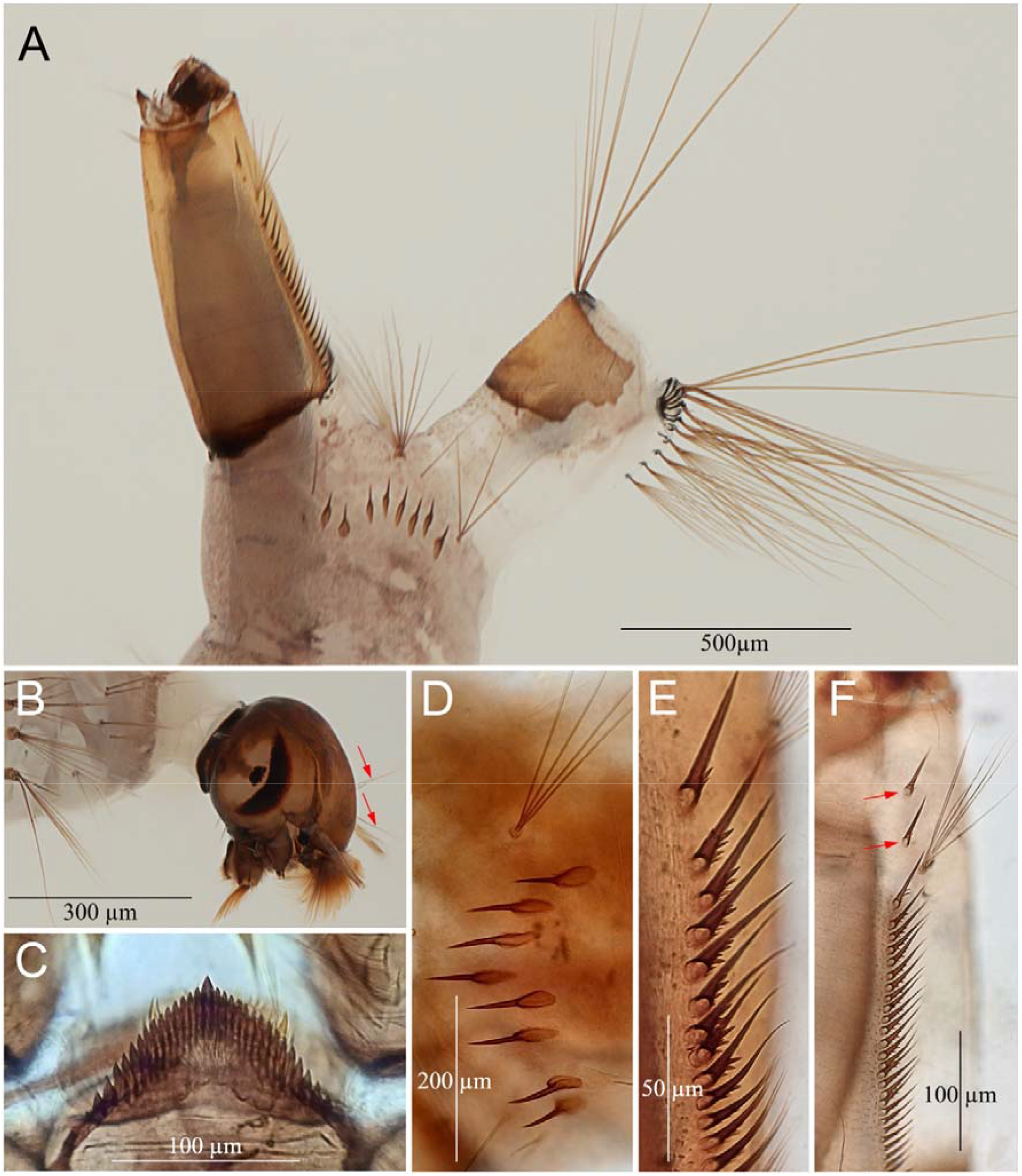
*Aedes vittatus*: morphological characteristic of the larval stage. A: Detail of abdominal segments VIII-X (the long anal papillae are missing); B: head showing single setae 5-C and 6-C (arrows); C: mentum; D: comb VIII with 8 spines; E: detail of the pecten teeth with 2 or 3 denticles; F: atypical pecten with the rare presence of 2 teeth (instead of one, arrows) inserted distal to the 1-S setae.

### Molecular identification

Morphological identification was confirmed with COI-based DNA barcoding. The *Ae. vittatus* specimens (Haiti_06, Haiti_07, Haiti_08, Haiti_09) cluster with Caribbean and American haplotypes (Cuba, Jamaica, the Dominican Republic, and Mexico) ^(3,5,24,25,26,27)^. In the haplotype network, they share closely related haplotypes separated by only a small number of mutations, suggesting a high level of genetic similarity within this group and particularly with Mexico^(5)^ (Figure 3). This Caribbean/American cluster appears distinct from African haplotypes and from most Asian haplotypes, which form separate groups within the network. The results support a common Asian-rather than African or European origin with regional spread in the Caribbean. Sequences were submitted to GenBank under accession numbers PZ344144, PZ344145, PZ344146 and PZ344147 and will be released upon publication.

**Figure 3.**
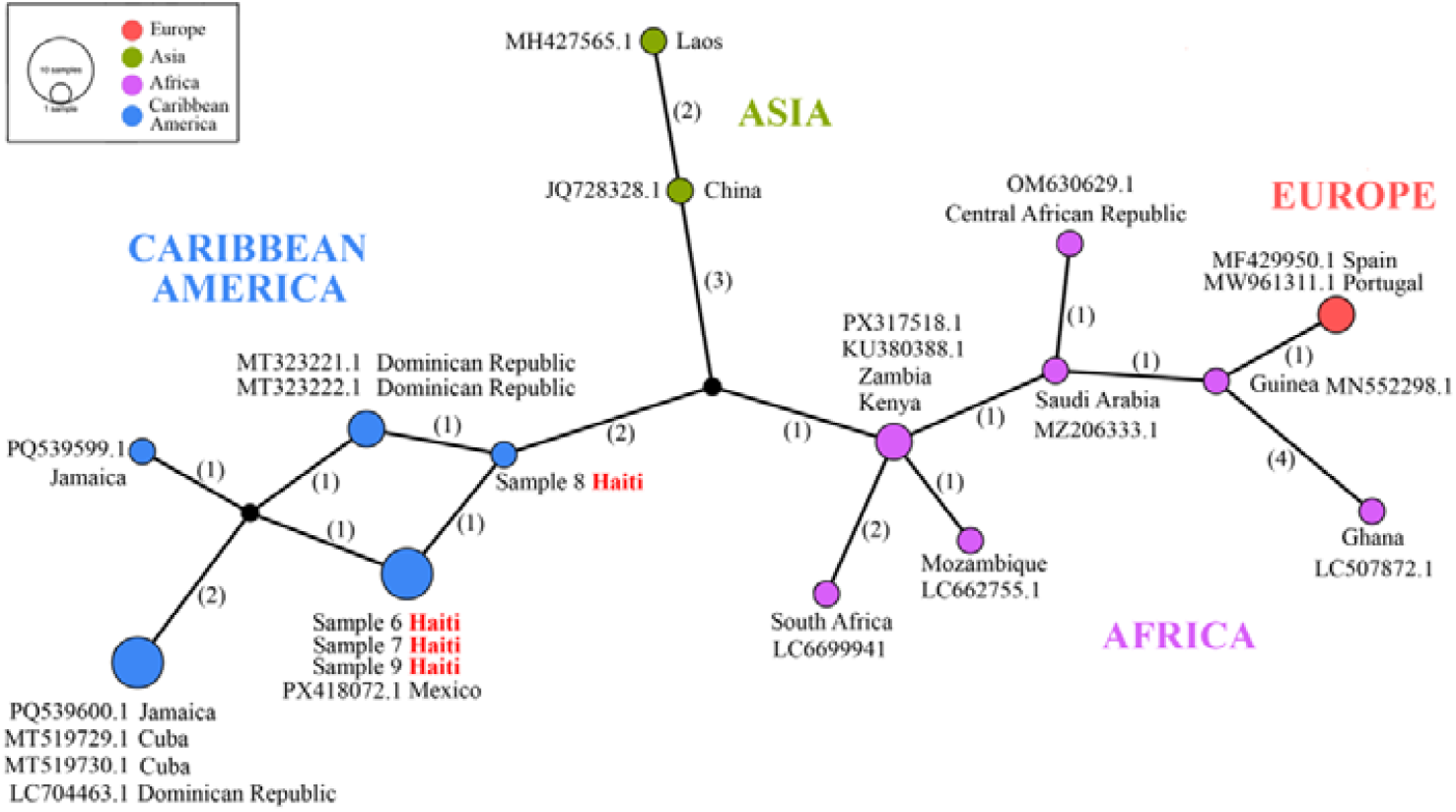
Haplotype network and phylogenetic tree resulting from the analysis of the COX I dataset. The haplotype network reconstruction considers missing data and gap. Values between parentheses on nodes indicate the inferred number of mutation steps between haplotypes or ancestral haplotypes (symbolized by a black node). The phylogenetic tree corresponds to the results of a Bayesian inference analysis (see text for details).

## Discussion

Species identification was confirmed morphologically and by COI-based DNA barcoding. Haitian *Ae. vittatus* specimens clustered with Caribbean and American haplotypes (Cuba, Jamaica, Dominican Republic, and Mexico) and were clearly distinct from African and most Asian lineages (Fig. 3). This pattern supports a shared regional origin and spread within the Caribbean rather than multiple independent introductions from the Old World.

Detection of *Ae. vittatus* in one-third of the sampled habitats and half of the surveyed communes, despite limited sampling, suggests that the species is already established and widespread in northern Haiti. Its absence from earlier surveys in 2013^(6)^, combined with its detection in the Dominican Republic in 2019^(3)^, supports a recent introduction to the island of Hispaniola, most likely facilitated by cross-border dispersal. Human-mediated transport probably contributed to its spread, particularly maritime trade and air travel. Passive transport via water-holding goods is a well-documented pathway for mosquito invasions, and the species’ desiccation-resistant eggs further enhance its dispersal potential. Genetic similarity with recently detected Mexican populations^(5)^ suggests strong regional connectivity, although the direction of spread remains unclear.

Current socio-environmental conditions in Haiti, including population displacement, informal settlements, inadequate water infrastructure, and unmanaged waste, favor the proliferation of container-breeding mosquitoes and likely facilitate the establishment of *Ae. vittatus*. The species exhibits high ecological plasticity, colonizing a wide range of artificial and natural habitats. Its tolerance to desiccation, salinity, and high temperatures further enhances its invasion potential^(28)^. Co-occurrence with *Ae. aegypti* and *Ae. albopictus* raises questions about interspecific interactions and future ecological dynamics in urban environments. Notably, we also reported *Cx. garciai* and *Cx. interrogator* from Haiti for the first time.

From a public health perspective, *Ae. vittatus* is competent for multiple arboviruses, including dengue, chikungunya, Zika, and yellow fever^(29,30)^. Its establishment adds complexity to Haiti’s epidemiologic context, which already includes endemic dengue, recurrent chikungunya outbreaks, and ongoing malaria and filariasis transmission. Notably, ecological niche models suggest that *Ae. vittatus* could find year-round suitable conditions across coastal regions of the Gulf of Mexico and extend into the southern Atlantic states of the United States, as well as large areas of Mexico and Central America^(31)^. This potential range expansion highlights the risk of further spread into North America, Central and South America and underscores the need for strengthened surveillance and preparedness beyond the Caribbean.

Our findings provide the first confirmed record of *Aedes vittatus* in Haiti and indicate that the species is likely established across multiple habitats and locations. Continued surveillance is needed to determine its distribution, ecological interactions, and potential contribution to arbovirus transmission in the region and beyond.

## Acknowledgments

We acknowledge funding from the Anténor-Firmin doctoral grant, the French Embassy in Haiti, the v, and the ARTS grant from the Institut de Recherche pour le Développement.

## Author’s contribution

DSL, PB, FS, DR – Conceptualization; DSL, PB, CL – Investigation; DSL, PB, LT, DR – Formal analysis; DR, FS, EE – project administration, supervision; DSL, FS, LT, DR – visualization; DSL, PB, DR - Writing – original draft, DSL, PB, LT, CL, EE, FS, DR - Writing – review & editing.

## References

1. Roiz D, Pontifes PA, Jourdain F, Diagne C, Leroy B, Vaissière AC, et al. The rising global economic costs of invasive Aedes mosquitoes and Aedes-borne diseases. Sci Total Env. 2024; 933:173054. 10.1016/j.scitotenv.2024.173054

2. Pabst R, Sousa CA, Essl F, Garcia-Rodriguez A, Liu D, Lenzner B, et al. Global invasion patterns and dynamics of disease vector mosquitoes. Nature Comm. 2025;16: 9127. 10.1038/s41467-025-64446-3.

3. Alarcón-Elbal P, Rodríguez-Sosa M, Newman B, Sutton W. The First Record of Aedes vittatus (Diptera: Culicidae) in the Dominican Republic: Public Health Implications of a Potential Invasive Mosquito Species in the Americas. Journal of medical entomology. 2020; 57(6): 2016□2021. 10.1093/jme/tjaa128.

4. Pagac BB, Spring AR, Stawicki JR, Dinh TL, Lura T, Kavanaugh MD, et al. Incursion and establishment of the Old World arbovirus vector Aedes (Fredwardsius) vittatus (Bigot, 1861) in the Americas. Acta Tropica. 2021; 213: 105739. 10.1016/j.actatropica.2020.105739

5. Chan-Chable RJ, Rodríguez-Luna C, Espinal-Palomino R, Ibarra-Cerdeña C. Detection of Aedes (Fredwardsius) vittatus Mosquitoes, Yucatán Peninsula, Mexico. Emerg Infect Dis. 2025; 31(11). 10.3201/eid3111.251358

6. Samson DM, Archer RS, Alimi TO, Arheart KK, Impoinvil DE, Oscar R, et al. New baseline environmental assessment of mosquito ecology in northern Haiti during increased urbanization. J Vector Ecol. 2015; 40(1): 46–58. doi:10.1111/jvec.12131.

7. ARIM [internet]. Arthropodes d’intérêt médical. Institut de Recherche pour le Développement (IRD). https://arim.ird.fr/arim/

8. Schaffner F, Angel G, Geoffroy B, Hervy JP, Rhaiem A, Brunhes J. Les moustiques d’Europe: logiciel d’identification et d’enseignement. CD-ROM. IRD. 2001; Editions, Paris, France.

9. Huang YM, Ward RA. A Pictorial Key for the Identification of the Mosquitoes Associated with Yellow Fever in Africa. Mosquito Systematics. 1981;13(2): 138–149.

10. Robert V, Ndiaye E H, Rahola N, Le Goff G, Boussès P, Diallo D, et al. Clés illustrées d’identification des moustiques du Burkina Faso, Cap-Vert, Gambie, Mali, Mauritanie, Niger, Sénégal et Tchad. IRD. 2022. 10.23708/fdi:010084866.

11. Huang YM. Medical entomology studies-VIII. Notes on the taxonomic status of Aedes vittatus (Diptera: Culicidae). Contrib. Am. Entomol. 1977; Inst. 14: 113–132.

12. Folmer O, Black M, Hoeh W, Lutz R, Vrijenhoek R. DNA primers for amplification of mitochondrial cytochrome c oxidase subunit I from diverse metazoan invertebrates. Mol Mar Biol Biotechnol. 1994; 3:294–9.

13. Camacho C, Coulouris G, Avagyan V, Ma N, Papadoulos J, Bealer K et al. BLAST+: architecture and applications. BMC Bioinformatics. 2009; 10, 421.

14. Katoh K, Rozewicki J, Yamada KD. MAFFT online service: multiple sequence alignment, interactive sequence choice and visualization. Brief Bioinform. 2019; 20:1160–6. 10.1093/bib/bbx108

15. Larsson A. AliView: a fast and lightweight alignment viewer and editor for large datasets. Bioinformatics. 2014; 30:3276–8. 10.1093/bioinformatics/btu531

16. Miller MA, Pfeiffer W, Schwartz T. Creating the CIPRES Science Gateway for inference of large phylogenetic trees. In: 2010 Gateway Computing Environments Workshop (GCE). New Orleans: IEEE; 2010. p. 1–8. 6.

17. Suchard MA, Lemey P, Baele G, Ayres DL, Drummond AJ & Rambaut A. Bayesian phylogenetic and phylodynamic data integration using BEAST 1.10. Virus Evol. 2018; vey016. 10.1093/ve/vey016

18. Rambaut A, Drummond AJ, Xie D, Baele G, Suchard MA. Posterior summarization in Bayesian phylogenetics using Tracer 1.7. Syst Biol. 2018; 67:901–4. 10.1093/sysbio/syy032

19. Rambaut A. FigTree. Edinburgh, UK: Institute of Evolutionary Biology, University of Edinburgh. 2018b.

20. Rozas J, Ferrer-Mata A, Sánchez-DelBarrio J C, Guirao-Rico S, Librado P, Ramos-Onsins S E, et al. DnaSP 6: DNA Sequence Polymorphism Analysis of Large Datasets. Mol. Biol. Evol. 2017; 34: 3299–3302. 10.1093/molbev/msx248

21. Bandelt HJ, Forster P, Röhl A. Median-joining networks for inferring intraspecific phylogenies. Mol Biol Evol. 1999; 16:37–48. 10.1093/oxfordjournals.molbev.a026036

22. Woolley SM, Posada D, Crandall KA. A comparison of phylogenetic network methods using computer simulation. PLoS One. 2008; 3: e1913. 10.1371/journal.pone.0001913

23. Leigh JW, Bryant D. popart: full-feature software for haplotype network construction. Methods Ecol Evol. 2015; 6:1110–6. 10.1111/2041-210X.12410

24. Díaz-Martínez I, Diéguez-Fernández L, Santana-Águila B, Atiénzar de la Paz E M, Ruiz-Domínguez D, Alarcón-Elbal PM (2021). Nueva introducción de Aedes vittatus (Diptera: Culicidae) en la región Oriental de Cuba: Caracterización ecológica y relevancia médica. InterAmerican Journal of Medicine and Health, 4(2021), 10–15. 10.31005/iajmh.v4i.175.

25. González, M. A., Bravo-Barriga, D., Rodríguez-Sosa, M. A., Rueda, J., Frontera, E., & Alarcón-Elbal, P. M. (2022). Species diversity, habitat distribution, and blood meal analysis of haematophagoushematophagous dipterans collected by CDC-UV light traps in the Dominican Republic. Pathogens, 11(7),): 714.

26. Noble SAA, Ali RL, Wilson-Clarke CF, Khouri NK, Norris DE, Sandiford SL (2025) Detection of invasive Aedes vittatus mosquitoes in Jamaica: molecular identification and surveillance implications. Parasites & Vectors, 18:469.

27. Tzuc-Dzul JC, Garcia-Rejon JE, Chi-Chim WA, Baak-Baak CM (2025) The first national record of the invasive mosquito Aedes vittatus (Diptera: Culicidae) in Mexico, a threat to public health in continental America.

28. Petersen V, Santana M, Karina-Costa M, Nachbar JJ, Martin-Martin I, Adelman ZN, et al. Aedes (Ochlerotatus) scapularis, Aedes japonicus japonicus, and Aedes (Fredwardsius) vittatus (Diptera: Culicidae): Three Neglected Mosquitoes with Potential Global Health Risks. Insects. 2024;15:600. 10.3390/insects15080600.

29. Sudeep AB, Shil P. Aedes vittatus (Bigot) mosquito: An emerging threat to public health. J Vector Borne Dis. 2017; 54: 295–300. 10.4103/0972-9062.225833

30. Diagne CT, Faye O, Guerbois M, Knight R, Diallo D, Faye O, et al. Vector Competence of Aedes aegypti and Aedes vittatus (Diptera: Culicidae) from Senegal and Cape Verde Archipelago for West African Lineages of Chikungunya Virus. Am J Trop Med Hyg. 2014; 91(3):635–641. 10.4269/ajtmh.13-0627

31. Ng’eno E, Nuñez-Penichet C, Ruiz-Utrilla ZP, Martins PI, Argudo V, Trindade WCF, et al. Potential distribution of Aedes vittatus as an invasive species in North America. PLoS One. 2025; 20(12): e0335534. 10.1371/journal.pone.0335534.

